# Free ISG15 as a dimer generates IL-1β-producing CD8α^+^ dendritic cells at the site of infection

**DOI:** 10.1101/111658

**Authors:** Anna Napolitano, Annemarthe G. van der Veen, Monique Bunyan, Annabel Borg, Svend Kjaer, Antje Beling, Klaus-Peter Knobeloch, Eva Frickel

## Abstract

ISG15 is strongly induced after type I IFN stimulation producing a protein comprised of two ubiquitin-like domains. Intracellularly, ISG15 can be covalently linked and modify the function of target proteins (ISGylation). In addition, free unconjugated ISG15 can be released from cells. We found that ISG15 is released in the serum of *Toxoplasma gondii* infected mice early after infection in a type-I IFN independent manner. Once in the extracellular space, free ISG15 forms dimers and enhances the release of key cytokines involved in the immune response to the parasite: IL-12, IFN-γ, and IL-1β. Its action is dependent on an actively invading and replicating live parasite. ISG15 induces an increase of IL-1β later during infection by leading to increased IL-1β producing CD8α+ dendritic cells at the site of infection. Here, we define for the first time the molecular determinants of active free ISG15 and link ISG15 to IL-1β production by CD8α+ dendritic cells. Thus we define ISG15 as a novel secreted modulator of the cytokine response during *Toxoplasma* infection.

---

ISG15 is a 15 kDa member of the family of interferon stimulated genes (ISGs) that contains two ubiquitin-like domains connected by a proline peptide linker. Similar to ubiquitin, ISG15 can be conjugated to intracellular host and viral proteinsemploying a cascade of E1 (UBE1L), E2 (UbcH8), and E3 conjugating enzymes. This ISGylation can modify protein function (LIT) and extensive studies have elucidated the role conjugated ISG15, mostly in the context of viral infection^1–3^. Recently the function of intracellular conjugated ISG15 has been extended to bacterial infections when it was found that *Listeria* infection induces ISGylation in non-phagocytic cells^4^. Interestingly, this ISG15 induction was type I IFN independent and lead to the release of IL-8 and IL-6 restricting bacterial replication^4^. Free ISG15 can also be released from the cell to the extracellular space and is detectable in the serum^5^

This extracellular free ISG15 it has long been appreciated to play an important role. *In vitro* studies suggested a role for free ISG15 as a cytokines able to induce IFN-γ secretion from NK cells^3^ and CD3+ lymphocytes *in vitro*^6,7^. Additionally, ISG15 was isolated from red blood cells of mice infected with *Plasmodium yoelii* and shown to act as a chemotractant for neutrophils^8^. However the *in vivo* function of free ISG15 remains ill-defined, and only two studies addressed the role offree ISG15 *in an in vivo* setting^9,10^. In a neonatal model of infection with Chikungunya virus, free ISG15 functions as an immunomodulator of proinflammatory cytokines, providing the first evidence that free ISG15 contributes to the host response during infection in the whole organism^9^. Most strikingly, it was recently demonstrated that ISG15 is a potent IFN-γ-inducing “cytokine” playing an essential role in anti-mycobacterial immunity^10^. Free extracellular ISG15 is effective alone or in synergy with IL-12 to induce IFN-γ secretion from granulocytes and NK cells in response to mycobacterial infection^10^. The ISG15-IFN-γ circuit may therefore be an “innate” complement to the more “adaptive” IL-12-IFN-γ circuit^10^.

*Toxoplasma gondii* is an obligate intracellular parasite that can infect virtually any nucleated cell. The active acute infection is believed to be mostly controlled by IFNγ, however, the parasite is never eliminated from an infected host and establishes a chronic infection at immune privileged sites, such as the brain and the heart^11^. In immunocompromised adults, this intracellular parasite dormant state in the brain reactivates and leads to the development of toxoplasmosis. The subsequent uncontrolled parasite replication causes life-threatening brain damage that is characterized by brain abscesses and necrotic areas. Two defence molecules, IL-12 and IFN-γ, orchestrate protective immunity in infected hosts^11^. The role of the IL-12-IFN-γ circuit during *Toxoplasma gondii* infection has been extensively characterised^11,12^.

*Toxoplasma gondii* infection triggers the release of a broad spectrum of molecules that can control the immune response to the parasite (LIT). In particular recent work has drawn much attention to the role of the inflammasome and IL-1β in the control of *Toxoplasma* infection^1–3,13–15^. Host protective immunity against *Toxoplasma* is highly dependent on the inflammasome sensors NLRP1 and NLRP3^4,15^. Moreover, the activation of the inflammasome is also dependent on the *Toxoplasma* strain dependent, with type II parasite being able to induce NLRPs activation and IL-1β release^4,14,15^.

Given the strong dependence of a *Toxoplasma* infection on IL-12 and IFN-γ, coupled with the parasite’s newly discovered ability to induce the inflammasome, we employed *Toxoplasma* infection to delineate the *in vivo* activity of free ISG15 in relation to IL-1β production. Here we shown that free extracellular ISG15 is produced during live *Toxoplasma* type II infection. Free extracellular ISG15, but not intracellular conjugated ISG15, enhances the generation of IL-12, IFN-γ and IL-1β during infection with live *Toxoplasma gondii* parasites. As a prerequisite for function, ISG15 had to form dimers. ISG15 production during *Toxoplasma* infection resulted in recruitment of IL-1β-producing CD8α^+^ dendritic cells (DCs) to the site of infection. These data demonstrate a novel role of ISG15 as an immunomodulatory molecule of IL-1β production within the context of the whole organism and extend the antipathogenic repertoire of ISG15 from viral and bacterial to protozoan infections.

## Results

### *Toxoplasma gondii* infection induces to the type I IFN-independent production and release of free ISG15 into the serum

To analyse if ISG15 is released during *Toxoplasma gondii* infection, we infected mice with *Toxoplasma* type II and monitored ISG15 levels in the serum during the acute phase of infection. We first determined the serum ISG15 levels by ELISA (Fig 1A). As early as day 2 post-infection (p.i.) the amount of ISG15 released was higher than in uninfected control mice (Fig. 1A) and its levels continue to rise until day 4 p.i. (Fig. 1A). To ascertain specificity of the commercial ELISA, we demonstrated that no signal is observed in the serum of *Toxoplasma*-infected ISG15^−/−^ mice (Fig. S1A). Likewise, recombinant ISG15 protein was detected in a dose-dependent manner (Fig. S1B). ISG15 may exist in the extracellular spaces as a monomeric or dimeric protein^16^. To analyse the quaternary structure of free ISG15 during *Toxoplasma* infection, we analysed the serum of infected mice by SDS-PAGE and immunoblotting under non-reducing conditions. We confirmed the release of ISG15 early upon infection and noticed the presence of a 30 kDa band that may represents dimeric ISG15 (Fig. 1B top). The protein band intensity from four experiments was quantified by ImageJ software and the immunoblot for the IgG heavy chain was used to normalize the levels of ISG15 in the serum (Fig. 1B bottom).

**Figure 1:**
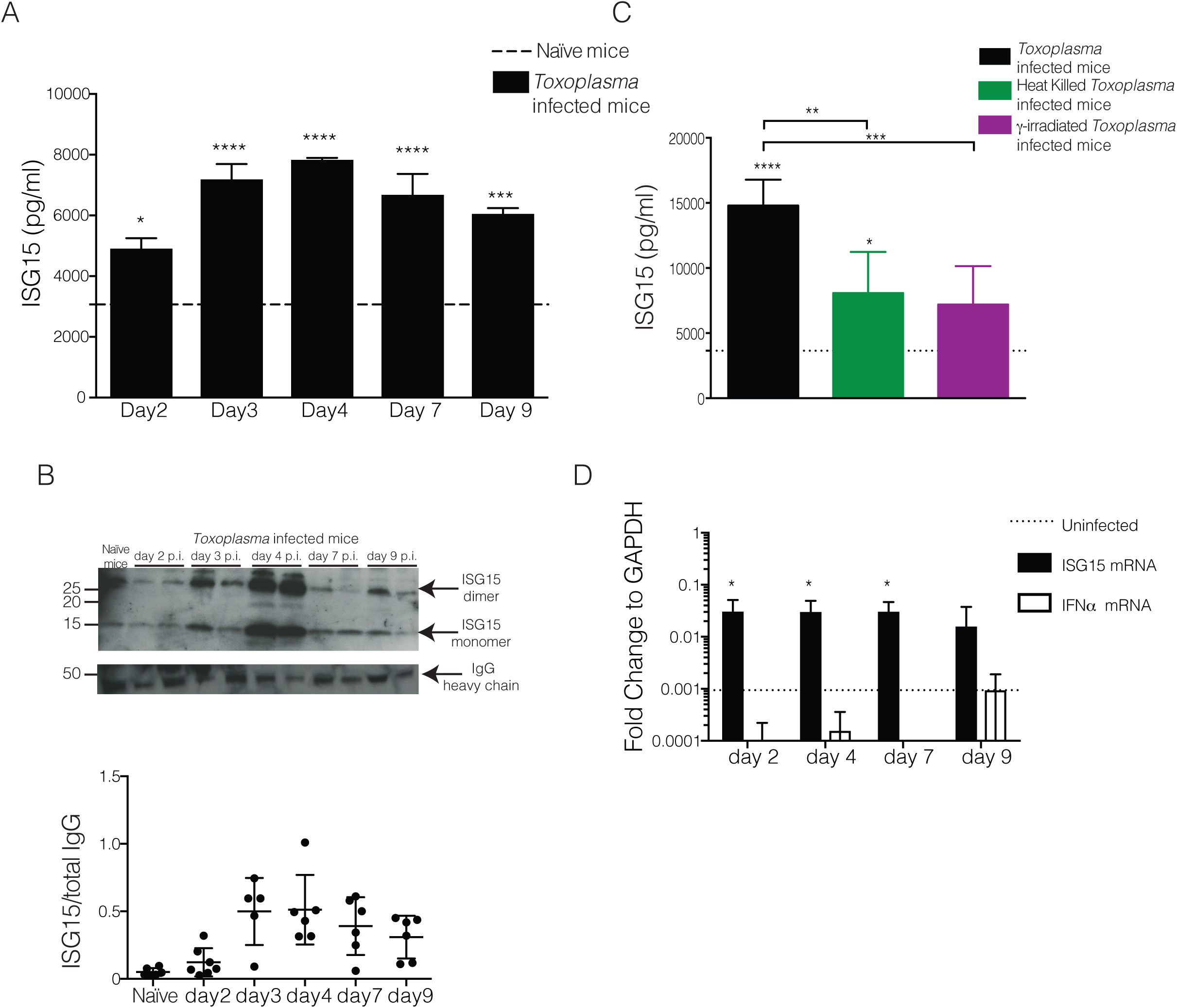
Extracellular free ISG15 levels increase upon Toxoplasma gondii infection in a type-I IFN independent way. C57BL6 mice were infected with 25,000 type II tachyzoites i.p. and serum samples were collected at different time points p.i.. A) Serum ISG15 levels during Toxoplasma infection measured by ELISA. 6 mice/group. One of three independent experiments. B) ISG15 immunoblot on serum samples. On the top, one representative blot of ISG15 and total IgG heavy chain used as loading control. On the bottom is plotted the quantification of four independent experiments. C) C57BL6 mice were infected with live or heat killed or γ-irradiated tachyzoites i.p., and serum ISG15 levels was assessed at day 4 p.i.. 3 mice/group, one of two independent experiments. D) qPCR for ISG15 and IFNα genes on RNA extracted from peritoneal exudate cells from uninfected and at day 2, 4, 7 and 9 p.i., 6 mice per group. One of two independent experiments. Statistics for all were analysed by two way ANOVA statistic with Tukey's multiple comparisons test, *p<0.05; ***p<0.0005; ****p<0.00005.

The release of ISG15 upon *Toxoplasma* infection might be an active process requiring with the presence of a live, replicative parasite, or be caused by specific recognition patterns of parasite or its products. To investigate whether ISG15 release depends on active host cell invasion by the parasite, we infected mice with either live parasites, γ-irradiated parasites that are able to invade but not replicate inside a cell, or heat killed parasites that are phagocytosed by cells. Only infection with live, actively replicating parasite leads to high ISG15 release in the serum was observed at day 4 p.i. (Fig. 1C), suggesting that ISG15 release during the early phase of infection is dependent on active invasion and replication of the parasite.

ISG15 belongs to the group of type I IFN inducible genes and is highly induced after type I IFN-generating viral infections^5,17 2,3,18^. However, it has recently been shown that ISG15 can also be produced during bacterial Listeriosis in a type I IFN-independent manner^6,7,19^. The role of type I IFN during protozoan parasite infection is controversial and not yet fully understood^20^. Recently a transcriptomic survey of the host response to different *Toxoplasma gondii* strains revealed that a subset of atypical strains induce a type I IFN response in macrophages and fibroblasts^8,21^. Other studies showed that *Toxoplasma* classical strains have the capacity to trigger a type I IFN response, but have evolved strategies to limit the induction of type I IFN and the ability of type I IFN to activate STAT1-dependent transcription^9,22,23^, highlighting new possible scenarios in the control of the parasitic infection. To determine if the increase of ISG15 during the early phase of *Toxoplasma* infection is caused by type I IFN, we analysed the expression of ISG15 and IFNα in peritoneal exudate cells at different time points post-infection and found that while there is an induction of ISG15 gene expression, IFNα expression is not induced by *Toxoplasma* infection (Fig. 1D). We concluded that early after infection *Toxoplasma* induces the release of ISG15 in a type-I IFN independent way. To further corroborate this finding we also analysed the expression of a panel of classical IFN target genes in the spleen of infected mice at day 4 p.i. (Fig. S1C). None of these genes were increased in expression upon *Toxoplasma* infection.

### Free extracellular ISG15 rather than ISGYlation enhances the release of IL-12, IFN-γ and IL-1β during infection with a live, replicative parasite

To dissect the role of free ISG15 during *Toxoplasma* infection, we infected mice with *Toxoplasma* type II and treated them with 1 μg of recombinant murine ISG15 at day 0, 1 and 2 p.i. by intra-peritoneal injection. As controls we employed both untreated *Toxoplasma* infected mice and naїve mice (Fig. S2A). We determined the levels of key cytokines released early after infection. Interestingly, shortly upon infection, ISG15-treated mice showed a modest but significant increase in IFN-γ, IL-12 and IL-1β, at different time points compared to infected but untreated mice (Fig. 2A). In particular, IL-1β secretion was consistently increased at all time points in ISG15-treated infected mice. Importantly, no increase in these cytokine levels were seen upon ISG15 treatment of uninfected mice, demonstrating that ISG15 alone is not sufficient to increase cytokine secretion, but only does so in the context of an infection (Fig. S2A). Additionally, this result ensures that the preparation of recombinant ISG15 is endotoxin free.

**Figure 2:**
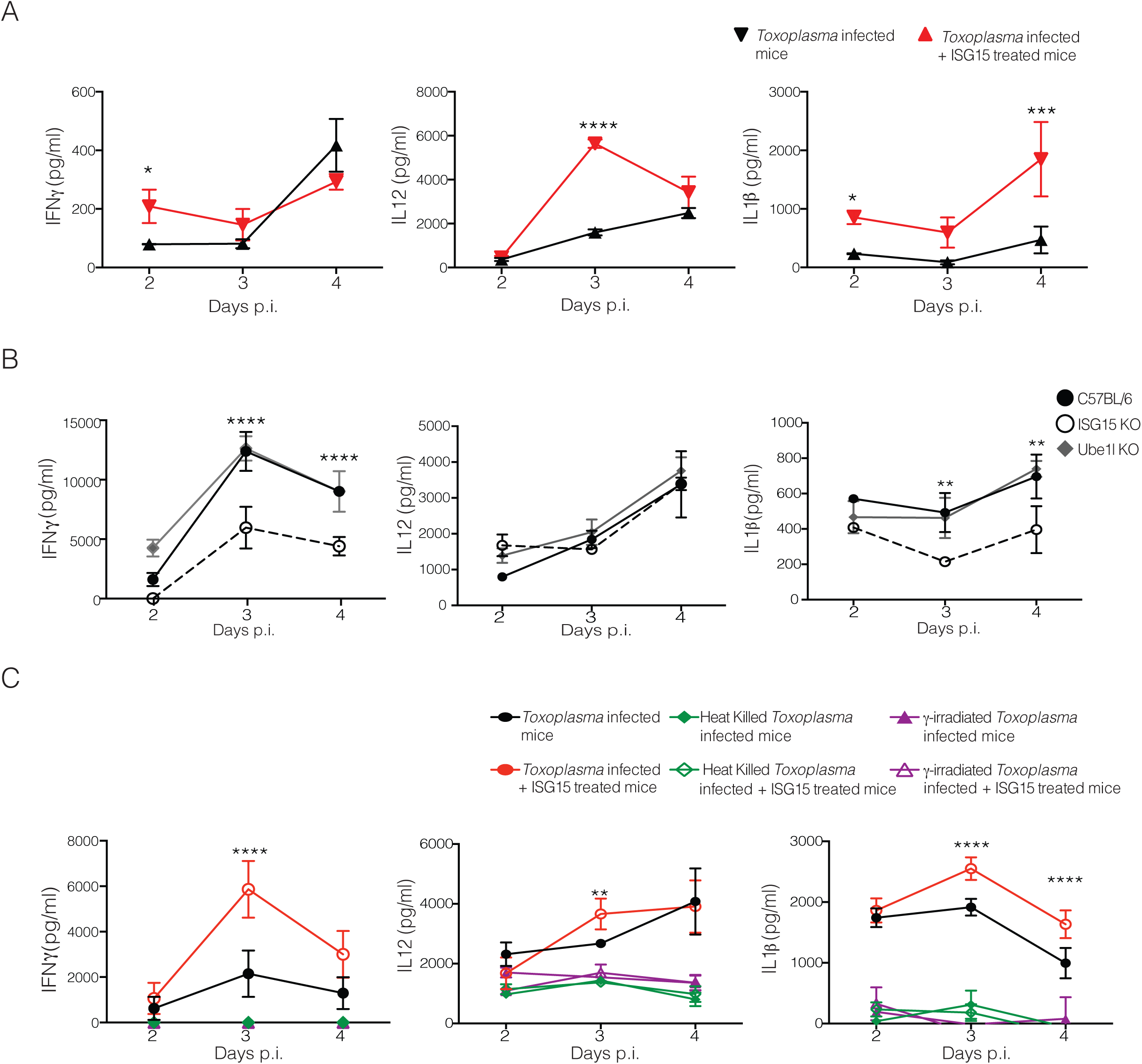
Extracellular ISG15 modulates IFN-γ, IL-12 and IL-Ιβ production during live *Toxoplasma* infection. A) C57BL6 mice were infected with 25,000 type II tachyzoites Pru. Mice were either infected only or infected and treated with recombinant ISG15 (1μg/mouse) at day 0, 1 and 2 post infection. ELISA for IFN-γ, IL-12 and IL-1β was performed on serum samples collected at different time points, 3 mice/group. One of three independent experiments. B) C57BL6, UbE1L−/− and ISG15−/− mice were infected with 25,000 type II tachyzoites i.p.. ELISA for IFN-γ, IL-12 and IL-1β was performed on serum samples collected at different time points, 3 mice/group. One of three independent experiments. C) Mice were infected with 25,000 type II live parasites, heat killed parasites or with _Y_-irradiated parasites. For each condition mice were either infected only or infected and treated with recombinant ISG15. ELISA for IFN-γ, IL-12 and IL-1β was performed on serum samples collected at different time points, 3 mice/group. One of two independent experiments. Statistics were analysed by two way ANOVA statistic with Tukey's multiple comparisons test, *p<0.05; **p<0.005; ***p<0.0005; ****p<0.00005.

As outlined in the introduction, only two studies have addressed the role of free extracellular ISG15 as a ‘cytokine-like’ molecule in an *in vivo* infection model ^9,10^ To asses whether the impact on cytokine secretion during *Toxoplasma* infection is attributable to the free or conjugated form of ISG15, we infected ISG15^−/−^ mice, that completely lack ISG15^11,24^, or UbE1L^−/−^ mice, which are devoid of intracellular conjugation of ISG15 through ISG15 E1 enzyme deficiency^11,25^. Upon infection levels of released cytokines were only diminished in ISG15^−/−^ but not UbE1L^−/−^ mice (Fig. 2B), strongly suggesting ISGylation is not critical for this process. However we did not observe a difference in survival or parasite load in the ISG15^−/−^ mice as compared to control mice (Fig. S2B and C). We therefore conclude that, albeit not necessary for survival, the free extracellular form of ISG15 modulates cytokine release during *Toxoplasma* infection.

To unequivocally assess if the cytokine modulatory function of ISG15 requires a process of active invasion from the parasite, we infected mice with live-, γ-irradiated-, or heat killed *Toxoplasma.* In each case we either only infected the mice or in addition treated the animals with recombinant ISG15. Recombinant ISG15 injection only increased cytokine release when mice were infected with live, actively replicating parasites (Fig. 3A), suggesting that ISG15-dependent modulation of cytokine levels during the early phase of infection is strictly dependent on active invasion and replication of the parasite.

**Figure 3:**
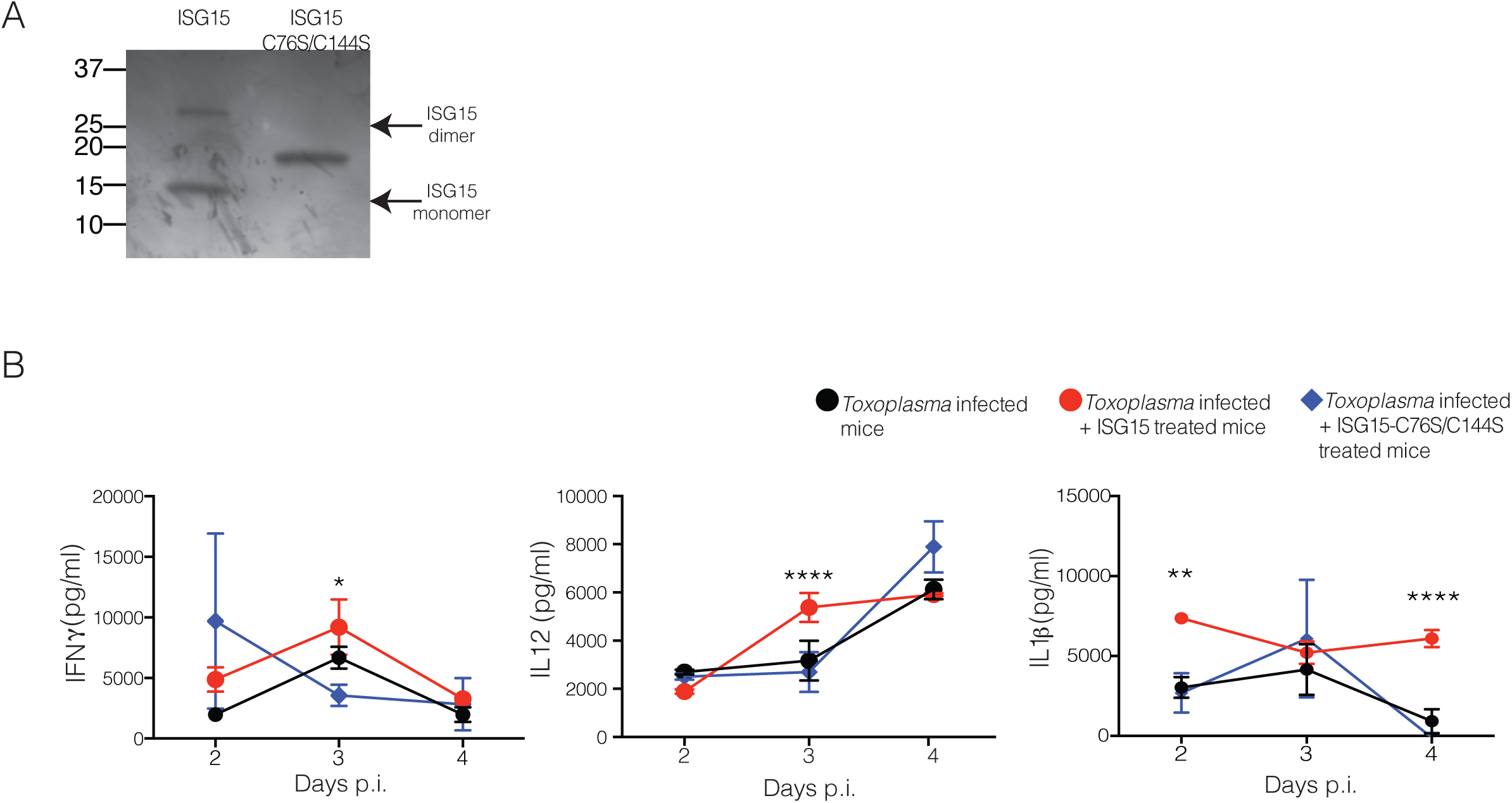
Free ISG15 forms dimers and induces cytokine release upon live Toxoplasma gondii parasite infection. A) Recombinat ISG15 and ISG15-C76S/C144S were left at RT for 1hr and then 1μg was run on SDS-PAGE with not reducing conditions and Comassie blue staining was performed on the gel. C) Mice were infected with live parasite and treated either with recombinant ISG15 (1μg/mouse) or with ISG15-C73A/C143A (1μg/mouse) at day 0, 1 and 2 p.i.. B) C57BL6 mice were infected with 25,000 type II tachyzoites Pru and were either infected only or infected and treated with recombinant ISG15 or ISG15-C76S/C144S (1μg/mouse). ELISA for IFN-γ, IL-12 and IL-1β was performed on serum samples collected at different time points, 3 mice/group. One of four independent experiments. Statistics were analysed by two way ANOVA statistic with Tukey's multiple comparisons test, *p<0.05; **p<0.005; ***p<0.0005; ****p<0.00005.

### Free ISG15 needs to dimerise to enhances the release of IL-12, IFN-γ and IL-1β during *Toxoplasma* infection

As we had noted the appearance of an anti-ISG15 immunoreactive 30 kDa band that may represent dimeric ISG15, we speculated that a dimeric form of ISG15 there is causative for the cytokine induction (Fig. 1B). As previously described, the disulphide bonds between cysteines in the hinge region might be responsible for ISG15 dimer formation^26^. To evaluate the role of ISG15 dimer formation on the modulation of cytokine release, we infected mice with *Toxoplasma* and left them untreated or treated them with recombinant ISG15, or ISG15 in which the two cysteines in the hinge region were replaced by alanine (ISG15 C76S/C144S). To show that the ISG15 C76S/C144S mutant cannot form dimers, we incubated ISG15 and its C76S/C144S mutant at RT for 1hour and then analysed the proteins under non-reducing conditions on an SDS-PAGE gel (Fig. 3B). When injected in infected mice, only the dimer-forming ISG15 has the capacity to increase the levels of cytokines in the serum (Fig. 3C). We therefore concluded that dimeric ISG15 modulates cytokine release during the early phase of a live *Toxoplasma* infection.

### Dimeric free ISG15 increases IL-1β-production by CD8α^+^ DCs at the site of infection

As many cell types have been identified as IL-1β-producers during infections^27,28^, we investigated which cells might release IL-1β in an ISG15 dependent manner during *Toxoplasma* infection. C57BL/6 mice were infected and left untreated or treated them with recombinant ISG15. At day 4 p.i., representing the peak of ISG15 release, we isolated cells from spleen, lymph nodes and peritoneal exudate and analysed the identity of cells of the innate cell compartment: neutrophils, macrophages, inflammatory monocytes and dendritic cells. In the spleen and the lymph nodes we did not find any notable difference in any cell population in PBS versus ISG15-treated mice (Fig. S3). However, at the site of infection, within the peritoneal exudate, we found an increased number and frequency of DCs in ISG15-treated infected mice (Fig. S4A). DCs play a critical role in the immune response to *Toxoplasma* infection^11,12,29^. Among the different subsets of DCs that are involved in the immune response to *Toxoplasma* infection, CD8α^+^ DCs play an important role and are involved in the presentation of *Toxoplasma*-specific epitopes to CD8^+^T cells^30^. We therefore analysed the DC compartment of the peritoneal exudate of Toxoplasma-infected ISG15^−/−^ and C57BL6 mice upon infection, the latter left untreated or treated with recombinant ISG15, or with ISG15- C76S/C144S. At day 4 p.i., we observed a higher number and frequency of both CD103^+^ and CD8α^+^ conventional DCs in infected mice treated with ISG15, whereas a significantly lower number and frequency of these cells was observed in the ISG15^−/−^ infected mice; finally mice treated with ISG15- C76S/C144S resemble the untreated mice, confirming that ISG15 has to form dimers to exert its immunomodulatory functions (Fig. 4A and S4B).

**Figure 4:**
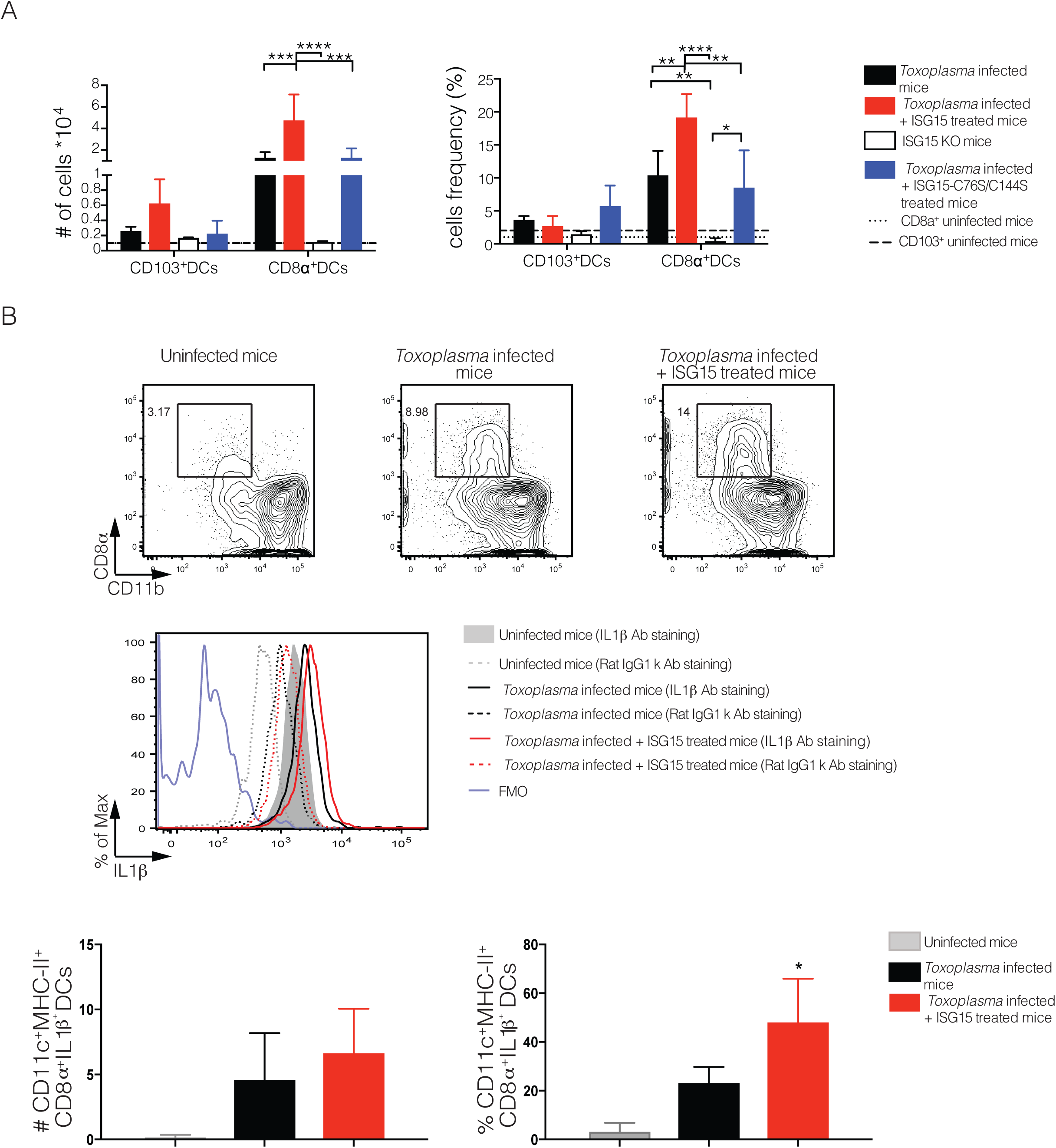
Free ISG15 recruits IL-1β-producing CD8α+ DCs to the site of infection. A) C57BL6 and ISG15−/− mice were infected with 25000 tachyzoite Pru i.p. . C57BL6 mice were either infected only or infected and treated with recombinant ISG15 or with ISG15-C76S/C144S mutant. At day 4 p.i. cells were isolated from the peritoneal exudate and FACS analysis was performed. Total cell numbers and cell frequency as percent of total is shown. 3 mice/group, one of four independent experiments. B) C57BL6 mice were either infected only or infected and treated with recombinant ISG15. At day 4 p.i. cells were isolated from the peritoneal exudate and CD8α+ population was enriched via MACS cell-sorting (negative purification). IL-1β intracellular staining was then performed on the enriched population. Top, one representative plot. Bottom, quantification of data combined from four independent experiments. Statistics were analysed by two way ANOVA statistic with Tukey's multiple comparisons test, *p<0.05; **p<0.005; ***p<0.0005; ****p<0.00005.

To determine whether the CD8α^+^ DCs present at the site of *Toxoplasma* infection are responsible for the increased IL-1β release in ISG15-treated mice, we performed an intracellular staining for IL-1β on these cells at day 4 p.i.. Mice infected and treated with ISG15 had an increased frequency and number of IL-1β-producing CD8α^+^ DCs (Fig. 4B). As control for the specificity of the IL-1β staining, we used both the fluourescence minus one (FMO) setting and an isotype-matched control antibody (Fig. 4B). These results suggest that early during *Toxoplasma gondii* infection ISG15 release contributes to IL-1β production by CD8α^+^ DCs present at the site of the infection.

## Discussion

We demonstrate that free extracellular ISG15 is involved in the regulation of the immune response to a protozoan infection with *Toxoplasma gondii in vivo*. We show that this free form of ISG15 is released in the serum early upon *Toxoplasma gondii* infection. Extracellular ISG15 functions as a dimer to modulate the release of IFN-γ, IL-12 and IL-1β. The enhanced production of IL1-β by ISG15 release is dependent on a live replicative parasite and is generated by CD8α^+^ DCs at the site of infection. Dimeric free ISG15 is therefore a yet undiscovered player in a balanced immune response to a protozoan infection.

The ubiquitin-like molecule ISG15 has been postulated to possess a cytokine-like role for more than 3 decades. Most studies have relied on *in vitro* stimulation with recombinant ISG15 of peripheral blood lymphocytes, purified CD3^+^T cells or CD56^+^ NK cells, or cell lines to show how this molecule could induce the release of IFN-γ^6 7^ Owhashi and colleagues, using a chemotaxis assay with different monocytes population isolated from mice peritoneal lavage, showed that ISG15 can recruit neutrophils^8^. While these are all intriguing studies, some doubts over the purity of recombinant ISG15 has remained, as well as the central question whether ISG15 *in vivo* can exert a cytokine-like function. Only two *in vivo* studies have been carried out assessing the role of ISG15 as an immunomodulatory molecule. It was first shown that ISG15 can regulate the host anti-viral immune response in a neonatal mouse model of Chikungunya virus infection^9,16^. Recently, Bogunovic and colleagues studied the first ISG15 deficient patients, which are characterized by Mendelian susceptibility to mycobacterial disease (MSMD). In the course of these experiments *M. tuberculosis* infection in mice was used to demonstrate that ISG15 plays a role as an IFN-γ-inducing secreted molecule essential for antimycobacterial immunity^5,10^. Following these *in vivo* studies, we found that free ISG15 has immunomodulatory functions during *Toxopasma gondii* infection, extending its role from viral and bacterial to protozoan infections.

ISG15 can be detected either intracellulary or extracellulary^5–7^ During the intracellular ISGylation process, ISG15 dimer formation, through conserved cysteine residues, reduces the amount of ISG15 that can be coupled with the target proteins, a process that can be prevented by nitrosylation of those cysteine residues on the ISG15 molecule^4,26^. Destinct from the ISGylation process, it is unclear whether the quaternary structure of extracellular ISG15 is essential for its function. Here, we show that the formation of ISG15 dimers trough conserved cysteines, is important for the cytokine-like activity of free ISG15 during *Toxoplasma gondii* infection. Thus, dimerisation of ISG15 is either required for *de novo* recognition by its putative receptor on target cells or for the downstream signalling response that ensues upon stimulation of its receptor. We unequivocally demonstrated that free ISG15 possesses an immunomodulatory capacity during a protozoan infection, and the protein has to be present as dimers in the extracellular space to exert this function. It will be interesting to determine whether this observation is restricted to the infection with *Toxoplasma* or whether this is a general feature of free ISG15 function *in vivo.*

Several cell types were suspected to represent targets for the immunomodulatory function of free ISG15. Among those are neutrophils^8^, NK cells^3,7,9^, and cells of the adaptive immune system, such as T cells^7,10^. We found that ISG15 levels correlate with an increased number and frequency of CD103^+^ and CD8α^+^ conventional dendritic cells at the site of infection. These two DC subsets are not identical but share a number of phenotypic characteristics, most likely derived from an immediate precursor that develops into either CD8α^+^ DCs or CD103^+^ DCs, depending on the tissue it seeds^11,31^. In the context of *Toxoplasma* infection different studies have shown that CD8α^+^ DC are a critical source of IL-12 during the acute phase of infection and relocate to the T cell area of the spleen to promote IFN-γ production by T cells^14,15 32,33^ Even thought CD8α^+^ DC are mainly defined as lymphoid resident-DCs, there are reports of migratory DCs expressing the CD8 marker presents in lower frequency in different compartments^19,34^. CD8α^+^ have been shown to participate in antigen presentation and T-cell priming for several intracellular pathogens including *Listeria monocytogenes*^11,12,35^ and *Salmonella* typhimurium^13–15, 36^, supporting a general role for CD8α^+^ DCs in the priming of immunity to intracellular pathogens. We newly show that those cells can release IL-1β to counteract the parasite. Moreover, based on our results, it is tempting to speculate that CD8α^+^ DCs express the ISG15 receptor and, in response to increasing ISG15 levels during *Toxoplasma* infection, migrate to the site of infection.

Beside IL-12, CD8α+ DCs can release many other cytokines^13–15,21–23,37–39^ even thought the full range that they might produce is still subject of investigation. Both intracellular signals, e.g. LPS or exogenous RNA/DNA, and extracellular signals, e.g. microbial products such as flagellin and toxins or *Toxoplasma* profilin, are able to trigger the activation of the inflammasome complex and the release of IL-1β. IL-1β can also be released in an inflammasome-independent way, with a number of non-caspase proteases involved in the cleavage of pro-IL-1β^40^. The fact that this molecule is released upon a broad variety of stimuli highlights the essential role played by IL-1β during the early phases of an immune response. In this study we show that *Toxoplasma* infection triggers the release of ISG15 that, in turn, stimulates IL-1β production by CD8α^+^ DCs at the site of infection. Free ISG15 may therefore be a novel modulator of the inflammasome response to an infection and it remains to be investigated if other inflammasome activating infections can be influenced by ISG15.

Although ISG15 functions as an immunomodulatory cytokine during the early phase of *Toxoplasma* infection, the unaltered survival rate of the infected ISG15^−/−^ mice shows that it is not a key player in counteracting the parasite. Perhaps ISG15 released upon an active invasion by *Toxoplasma* might be a starting signal used by the invaded cells to initiate an immune response to the parasitic infection. In fact, the release of ISG15 is not the response to a general DAMP signal, but needs the parasite to actively enter into the cell, replicate and initiate the inflammatory response. Only a live, replicative parasite is able to induce such an immune response suggesting that ISG15 is freed as consequence of an active infection to work as a danger signal. Accordingly, while ISG15 alone is not sufficient to lower the parasite load, it can increase IL-1β levels.

In our study we found that free dimeric ISG15 triggers an increase in CD8α^+^ DCs producing IL-1β, associated with the immune response to the parasite *Toxoplasma gondii.* During the early phase of the immune response to *Toxoplasma* infection, ISG15 act as modulating cytokine released at the site of the infection. ISG15 is thus a new molecule involved in the immune response to *Toxoplasma* infection specifically and acts to enhance IL-1β production during this infection. Free dimeric ISG15 as an enhancer of the inflammasome response during infections may be a more broad phenomenon and warrants further investigation.

## Methods

### Mice

C57BL/6 and ISG15 deficient^7,24^ mice were housed and bred at the Francis Crick Institute Ltd, Mill Hill Laboratory under specific pathogen-free conditions. Ube1L-deficient mice^8,25^ were a kind donation of Antje Beling. Experiments were performed on 6- to 8-week-old males. All procedures involving mice were approved by the local ethical committee of the Francis Crick Institute Ltd, Mill Hill Laboratory and are part of a project license approved by the Home Office, UK, under the Animals (Scientific Procedures) Act 1986.

### Parasite culture and infections

*Toxoplasma* avirulent type II strain Pru (kind gifts from Marc-Jan Gubbels and Jeroen Saeij)^41 42^ were used for all the experiments. *Toxoplasma gondii* culture were maintained by serial passage on monolayers of human foreskin fibroblasts (HFFs) as described previously^43^. Freshly egressed parasites were harvested from HFF culture, filtered, counted, resuspended in PBS and injected intraperitoneally into mice. For *in vivo* imaging *Toxoplasma* expressing the GFP and firefly luciferase^41^ was used. Mice were injected i.p. with 3 mg firefly D-luciferin (Perkin Elmer), left for 10 min and imaged with an IVIS Spectrum-bioluminescent and fluorescent imaging system (Xenogen Corporation) under isoflurane anaesthesia (Abbott).

For *in vivo* experiments infected mice were treated with ISG15 (1 μg/mouse) or treated with either PBS, buffer from the gel filtration or nothing with no difference observed.

### Reagents and Antibodies

Anti-ISG15 rabbit sera, gently gift of Peter Knobeloch, was used as primary antibodies for immunoblot. Secondary antibodies for immunoblotting were from KCL: goat anti-rabbit HRP (474-1506) and goat anti-mouse HRP (474-1806), used for the IgG heavy chain blot. Fluorescently labeled antibodies for cytofluorymetry (FACS) against Ly6C (clone HK1.4), CD11b (clone M1/70), I-A/I-E (clone M5/114.15.2), CD11c (clone N418), Ly6G (clone 1A8), F4/80 (clone BM8), CD103 (clone 2E7), CD8α (clone 53-6.7), CD45R/B220 (clone RA3-6B2), CD49b (clone DX5), TCRβ (clone H57-597) and purified CD16/32 (anti Fc-g receptor, clone 93) were purchased from Biolegend. Anti-mouse IL-1β (clone NJTEN3) and Rat IgG1 k isotype control (clone eBRG1) were purchased from eBioscience. Anti-FITC MACS beads (130-048-701) used for negative selection for the intracellular staining were from Miltenyi Biotec. ELISA Pierce TMB substrate kit (34021) was from Thermo Fisher Scientific. Dulbecco’s PBS was from VWR (21-031-CV)

### Immunoblot and Comassie Blue staining

Immunoblot was used to quantitatively detect specific protein bands after separation of proteins on pre-casted 4-20% gradient SDS-PAGE gel (4561095) from Biorad. Proteins were transferred onto nitrocellulose membrane (IB301002, Life Technologies) by dry-blotting (iBlot®, Life Technologies). Blots were blocked at RT 1h in a 5% non-fat dried milk/PBS solution. The membrane was then incubated with primary antibody in 0.5% non-fat dried milk/PBS solution overnight at 4°C. On the next day, blots were washed five times for 20 min in PBS +0.1% Tween 20. Blots were incubated for 1h with HRP-conjugated secondary antibodies in Blotto+0.1% Tween 20. Blots were washed five times for 20 min in PBS +0.1% Tween 20. Finally, blots were developed with Immobilon Western Chemiluminescent HRP Substrate (Merck Millipore). Blots were exposed to CL-XPosure film (Thermo Scientific) and films developed. Membrane was then treated with Western blot Stripping Buffer (Thermo Fisher Scientific, 62299) for 10 minutes RT and blocked again at RT 1h in a 5% non-fat dried milk/PBS solution. After blocking membrane was incubate with the anti-mouse HRP secondary only for 1h at RT and developed as before. Comassie Blue staining was used to detect specific protein bands after separation of proteins on pre-casted 4-20% gradient gel. After the run gel was stained with Comassie Brilliant Blue R-250 (161-0436) from Biorad and then destained with water until bands were clear.

#### Flow cytometry

Single-cell suspensions were prepared from spleen and lymph nodes (LNs) by mechanical disruption. Spleens were treated with the Red Blood Cell Lysing buffer (Sigma, RNBD7167) for 5 minutes at room temperature to remove most tof the blood cells. Peritoneal exudate cells were harvested by peritoneum lavage with 5 mL of PBS. After counting, cells were resuspended in PBS 1% BSA (PBA) and stained for 20 min at 4°C in an appropriate antibody cocktail. Cells were washed twice with PBA before to read them at the FACS.

For the intracellular staining, after harvesting, cells were stained with a cocktail of FITC-antibodies and negatively selected with the anti-FITC beads according to the manufacturer’s instructions. Cells were then stained with the surface antibodies in PBA and BD Pharmigen Cytofix/Cytoperm kit (554714) was used for the intracellular staining according to the manufacturer’s instructions. Cells were run on a BD LSRII or BD Fortessa X20 and analyzed using FlowJo software (Tree Star).

### Quantification of cytokines by ELISA

Mice were bleeding at different time point according to the experiment. Blood was left at RT for a minimum of 20 minutes and serum was then isolated by centrifugation and stored at −80. Cytokines were analysed by enzyme-linked immunosorbent assay (ELISA). Commercially available kits were used according to the manufacturer’s instructions to quantify the concentration of ISG15 (Caltag Medsystem, CY-8091) IL-1β (BD Pharmigen, 559603), IFN-γ (BD Pharmigen, 555138) and IL-12/IL-23 p40 (R&D Sytems Inc., DY2398).

### Real Time PCR

RNA was extracted using Trizol reagent (Thermo Fisher Scientific, 10296010) and quantified at the Nanodrop 3300 (Thermo Fisher Scientific). RNA was then reverse transcribed with the Superscript cDNA Vilo Synthesis kit (Invitrogen, 11754050) according to the manufacturer’s instructions. TaqMan gene expression master mix (Applied Biosystems, 4369016) was used to perform Real Time PCR in triplicate with 100 ng cDNA/reaction using specific primers and Gadph (Applied Biosystems, Mm99999915_g1) endogenous control on ABI Prism 7900 (Applied Biosystems) and analysed with SDS 2.2.1 software. Relative quantitation of gene expression was determined using the comparative cycle threshold method^44^. The specific primers used were all from Applied Biosystem: Ifit3 (Mm01704846_s1); Oas1a (Mm00836412_m1); Oas2 (Mm00460961_m1); Mx1 (Mm00487796_m1); Mx2 (Mm00488995_m1); Tnfsf10 (Mm01283606_m1); Ifna2 (Mm00833961_s1); Isg15 (Mm01705338_s1).

### Statistical analysis

All statistical significance analyses were performed using Prism software (GraphPad Software). Comparisons of data were performed with using two ways ANOVA statistic with Tukey's multiple comparisons test. Survival rates were compared by log-rank survival analysis of Kaplan-Meier curves.

#### Purification of recombinant ISG15

Murine mature ISG15 fused to an HA tag, a TEV protease cleavage site, and a linker containing two glycine residues was cloned into the pTriEx6 expression plasmid. Quich change mutagenesis (Strategene) was used to introduce the C76S and C144S mutations. pTriEx6 containing the His-GST-3C-HA-TEV-GG-ISG15 gene was transfected into insect cells (Sf21 Berger) using the FlashBac system from Oxford Expression Technologies and Fugene HD as transfection reagent. 2.5 μl of FlashBac DNA, 5 μl of Fugene HD and 500 ng of pTriEx6 was used to transfect 4x105 cells/ml in a 6-well plate. Five hours post-transfection, 1 ml of Sf900 III containing fungizone was added to the wells. The cells were incubated for 5 days at 27°C. The resulting 2 ml P1 virus was used to infect 25 ml of Sf21 cells at 106 cells/ml for 3 days at 27°C. The resulting P2 virus was titered using qPCR and amplified to 50ml of P3 at a MOI of 1 (cells at 106/ml). The P3 virus was used to infect a large scale of Sf21 cells, either at low density (cells at 106/ml, MOI 1) or high density (cells at 6.106/ml, MOI 3, Glucose, Lactalbumine and Yeastolate added to the culture). The infected cells were pelleted and frozen at −80°C until purification. The intracellular ISG15 was purified from the Sf21 cell pellets using Glutathione Agarose (Cube Biotech) and Superdex 75 10/30 GL (GE Healthcare). Cells were lysed in 25 mM Tris pH7.5, 75 mM NaCl, 2.5% Glycerol, 0.5% Triton, Benzonase nuclease (Sigma) and protease inhibitor tablets (Roche) using a sonication probe on ice. The lysate was then spun at 20.000 g for 10 min at 4°C to pellet cell debris. The cleared lysate was incubated with Glutathione Agarose at 4°C for 2h. The beads were washed 3 times with 25 mM Tris pH7.5, 75 mM NaCl, 2.5% Glycerol. The protein was cleaved using TEV protease overnight at 4°C. The cleaved protein was further purified by gel filtration in 25 mM Tris pH7.5, 75 mM NaCl, 2.5% Glycerol. The fractions containing pure ISG15 were pooled and concentrated. SDS PAGE followed by Coomassie staining or immunoblotting with an anti-ISG15 antibody (Santa Cruz, clone A5, sc-166712) were used to verify purity of the recombinant product.

## Acknowledgements

We would like to thank the Frickel lab, Julie Helft, Sumana Sanyal and Caetano Reis e Sousa for productive advice and discussion. We thank the Crick FACS facility and Biological Services for expert technical assistance. This work was supported by the Francis Crick Institute, which receives its core funding from Cancer Research UK (FC001076), the UK Medical Research Council (FC001076), and the Wellcome Trust (FC001076). Eva-Maria Frickel was supported by a Wellcome Trust Career Development Fellowship (091664/B/10/Z).

## Author contribution

AN and EMF contributed to the intellectual design of the study. AN, AGV, MB conducted research. AN, AGV, AB, KPK wrote and edited the manuscript. AB and SK prepared recombinant ISG15. AB kindly donated Ube1L knockout mice. KPK kindly donated ISG15 knockout mice and reagents. EMF supervised the study.

---

**Figure S1:**
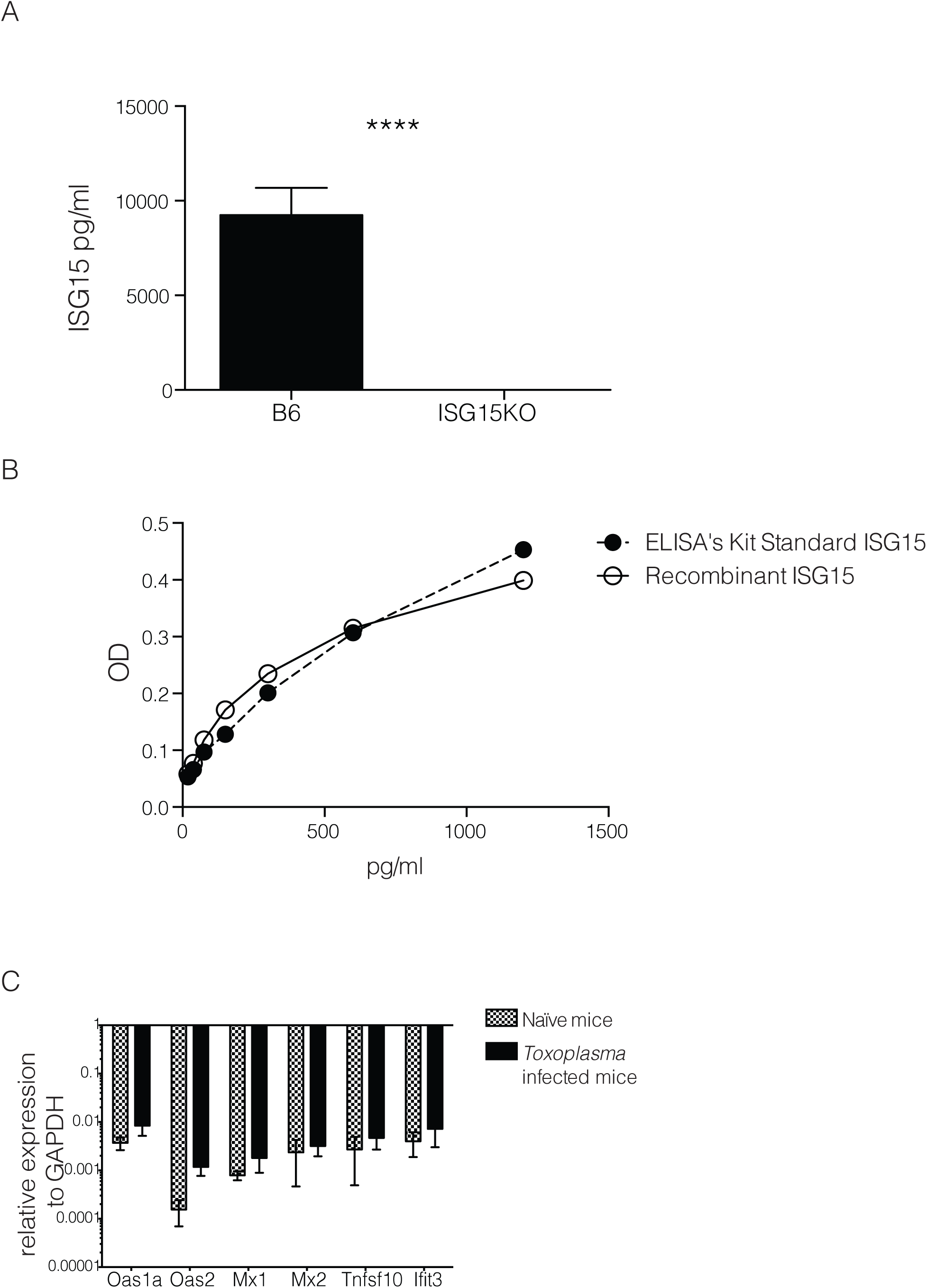
A) ISG15 ELISA on serum collected at day 4 p.i. from C57BL/6 and ISG15−/− mice infected with 25,000 type II tachyzoites i.p.. B) In house-made recombinant ISG15 was used as standard in comparison with the standard provided from the ELISA kit. C) qPCR genes activated by type-I IFN stimulation on RNA extracted from spleen of uninfected and at day 4 p.i., 6 mice per group. One of two independent experiments. Statistics for all were analysed by two way ANOVA statistic with Tukey's multiple comparisons test, ****p<0.00005.

**Figure S2:**
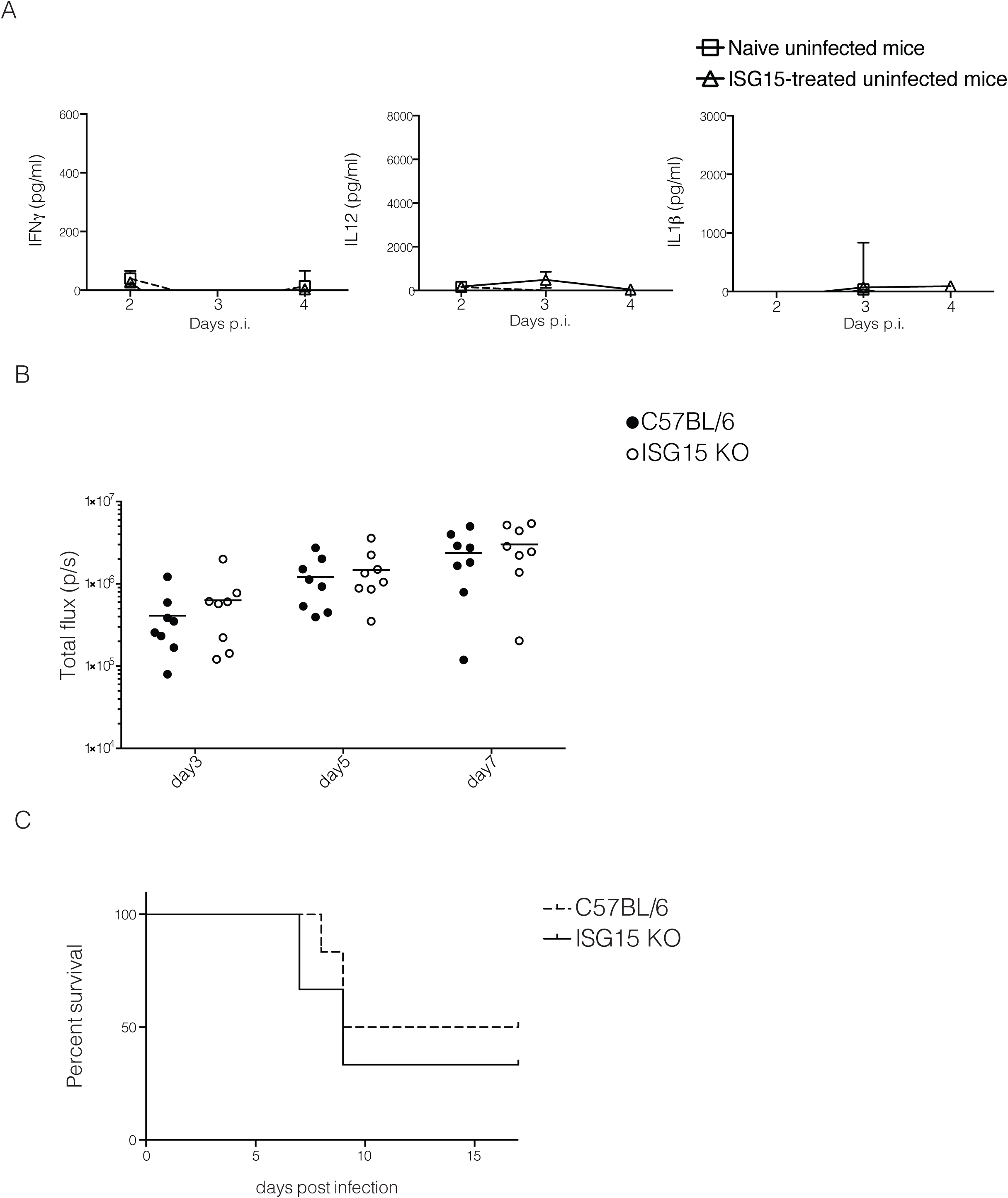
A) C57BL6 mice were left uninfected or treated with recombinant ISG15 (1μg/mouse) at day 0, 1 and 2 post infection. ELISA for IFN-γ, IL-12 and IL-1β was performed on serum samples collected at different time points, 3 mice/group. One of three independent experiments. B) and C) C57BL6 and ISG15−/− mice were infected with 25000 tachyzoite Pru i.p.. B) Parasite load was measured by in vivo imaging of the firefly luciferase signal. C) Survival of mice. Data combined from three independent experiments, 20 mice total.

**Figure S3:**
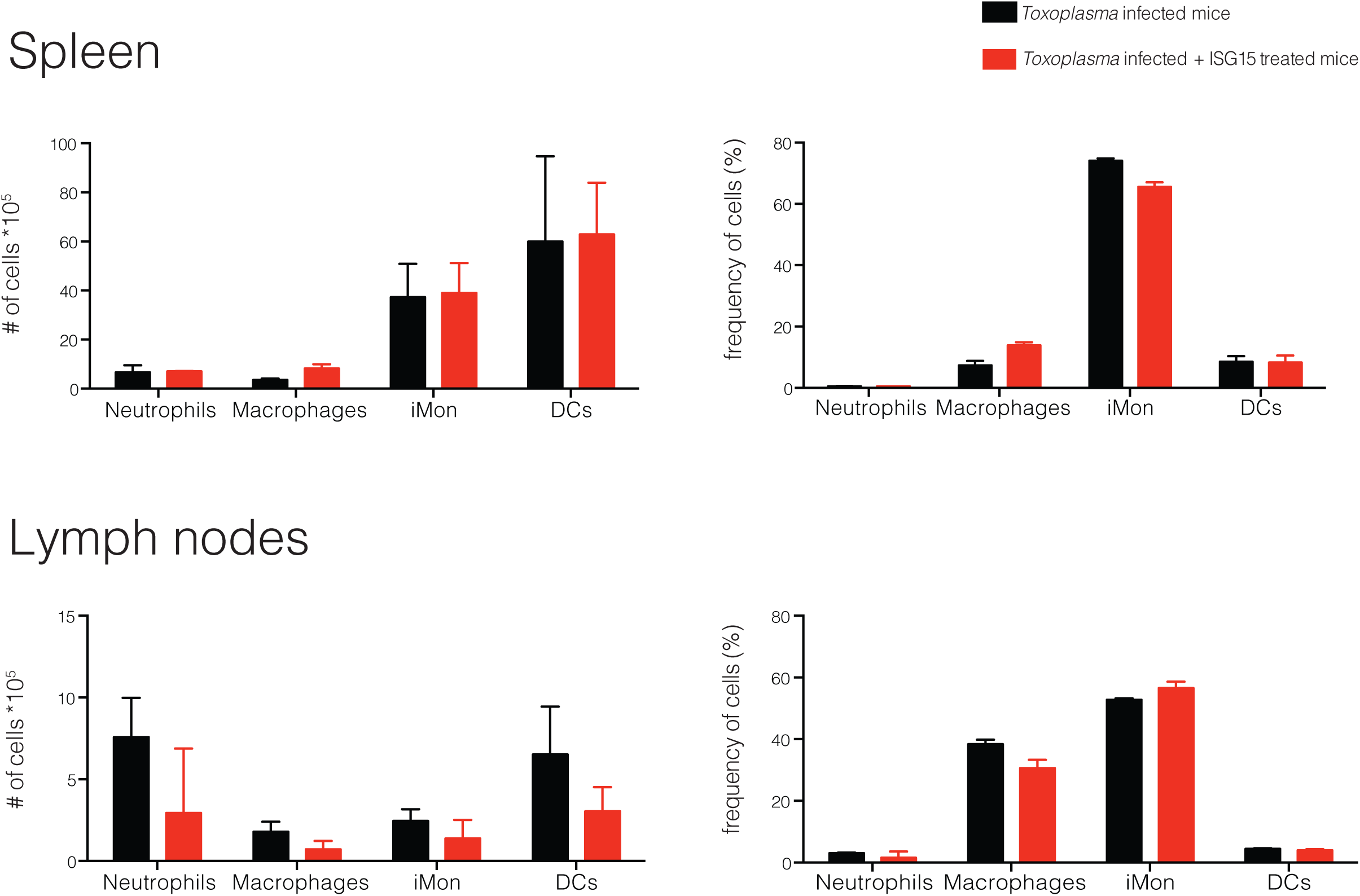
C57BL6 mice were either infected with 25,000 type II tachyzoites i.p. and treated with recombinant ISG15 (1μg/mouse) at day 0, 1 and 2 post infection or infected only. Mice were sacrified 4 days p.i. and cells were purified from draining lymph nodes (inguinal and brachial) and spleen. Number and frequency of the cell populations analysed is shown, 3 mice per group. One of three independent experiments.

**Figure S4:**
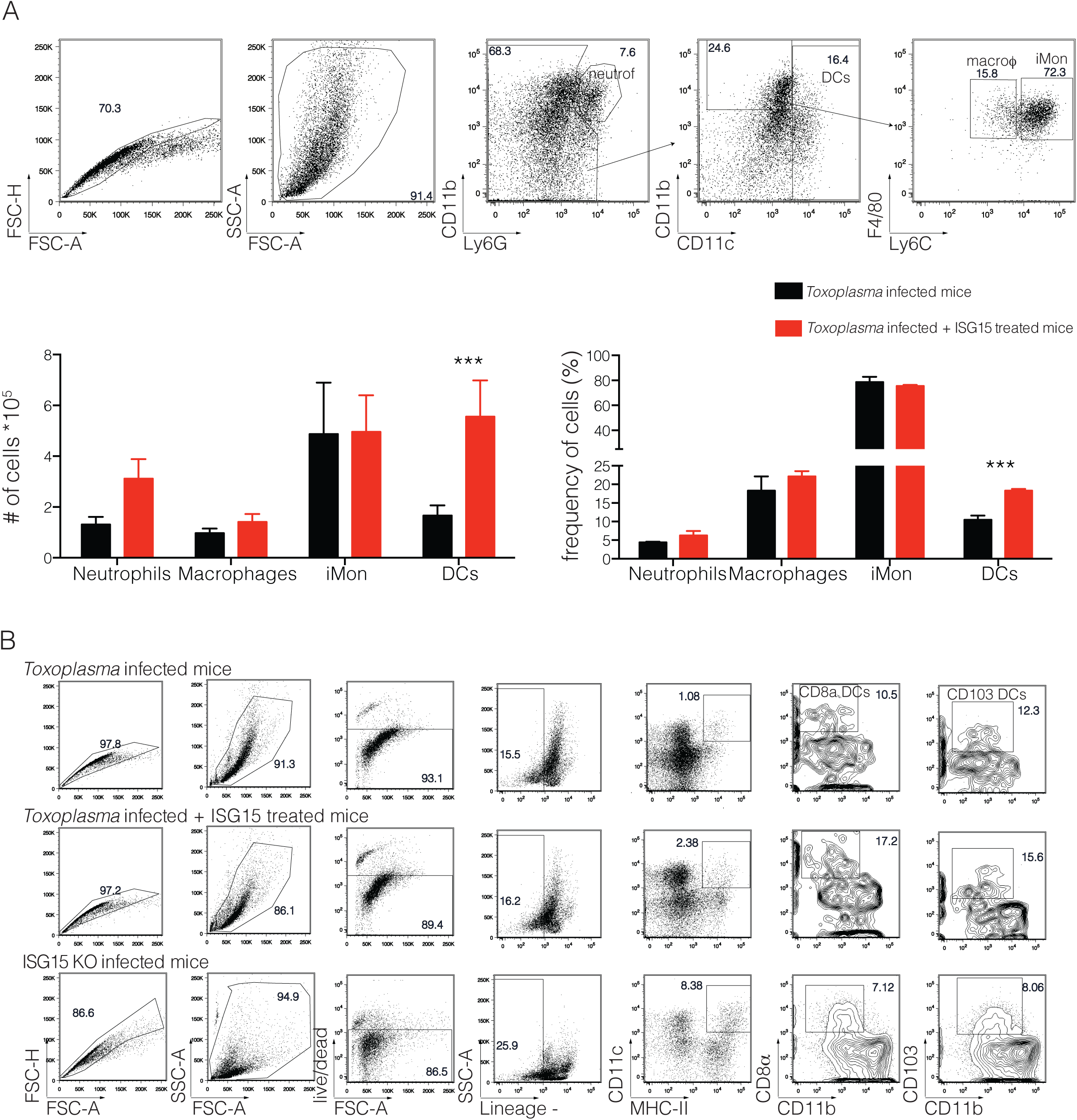
C57BL6 mice were either infected with 25,000 type II tachyzoite i.p. and treated with recombinant ISG15 or infected only. Mice were sacrified 4 days post infection and cells were purified from peritoneal exudate. A) Gating strategy (top) and quantification (bottom) for the staining used to analyse the innate cell populations, 3 mice per group. One of three independent experiments. B) Gating strategy for the staining used to analyse CD8α+ and CD103 DCs, 3 mice per group. One of three independent experiments. Statistics were analysed by two way ANOVA statistic with Tukey's multiple comparisons test, ***p<0.0005.

